# Gene Expression Changes in Cultured Reactive Rat Astrocyte Models and Comparison to Device-Associated Effects in the Brain

**DOI:** 10.1101/2023.01.06.522870

**Authors:** Ti’Air E. Riggins, Quentin A. Whitsitt, Akash Saxena, Emani Hunter, Bradley Hunt, Cort H. Thompson, Michael G. Moore, Erin K. Purcell

## Abstract

Implanted microelectrode arrays hold immense therapeutic potential for many neurodegenerative diseases. However, a foreign body response limits long-term device performance. Recent literature supports the role of astrocytes in the response to damage to the central nervous system (CNS) and suggests that reactive astrocytes exist on a spectrum of phenotypes, from beneficial to neurotoxic. The goal of our study was to gain insight into the subtypes of reactive astrocytes responding to electrodes implanted in the brain. In this study, we tested the transcriptomic profile of two reactive astrocyte culture models (cytokine cocktail or lipopolysaccharide, LPS) utilizing RNA sequencing, which we then compared to differential gene expression surrounding devices inserted into rat motor cortex via spatial transcriptomics. We interpreted changes in the genetic expression of the culture models to that of 24 hour, 1 week and 6 week rat tissue samples at multiple distances radiating from the injury site. We found overlapping expression of up to ∼250 genes between *in vitro* models and *in vivo* effects, depending on duration of implantation. Cytokine-induced cells shared more genes in common with chronically implanted tissue (≥1 week) in comparison to LPS-exposed cells. We revealed localized expression of a subset of these intersecting genes (e.g., *Serping1, Chi3l1, and Cyp7b1)* in regions of device-encapsulating, glial fibrillary acidic protein (GFAP)-expressing astrocytes identified with immunohistochemistry. We applied a factorization approach to assess the strength of the relationship between reactivity markers and the spatial distribution of GFAP-expressing astrocytes *in vivo*. We also provide lists of hundreds of differentially expressed genes between reactive culture models and untreated controls, and we observed 311 shared genes between the cytokine induced model and the LPS-reaction induced control model. Our results show that comparisons of reactive astrocyte culture models with spatial transcriptomics data can reveal new biomarkers of the foreign body response to implantable neurotechnology. These comparisons also provide a strategy to assess the development of *in vitro* models of the tissue response to implanted electrodes.

## Introduction

Lost nervous system function can be partially returned in tetraplegic patients with implanted microelectrode arrays (MEAs), as demonstrated in clinical trials^1–4^. This technology has helped these patients to communicate with external assistive devices to regain independence in their daily living. Likewise, implanted electrodes have the potential to treat epileptic, Alzheimer’s disease, and Parkinson’s disease patients, as well as patients suffering from substance use disorders. Although the potential impact of this technology is profound, an ongoing challenge is presented by the instability and limited longevity of recorded signals detected by implantable electrodes. It has long been suspected that the foreign body response (FBR) to the implant contributes to the eventual loss of signal^5–7^. Researchers have responded by creating a myriad of next generation probe designs to mitigate the tissue response, and each new design comes with unique benefits and limitations^8^. Understanding the biological response to electrodes may offer an avenue to improve chronic performance, understand the relationship between performance and design features, and amplify the already-compelling therapeutic promise of these devices.

The stereotypical pathophysiologic response to MEAs occurs in the following stages: (1) device insertion typically causes mechanical damage to tissue, breach of the blood-brain barrier (BBB) and disruption of vasculature at the implantation site; (2) microglia are activated to encapsulate the probe immediately thereafter, creating a physical barrier responsible for limiting ionic exchange with the probe while potentially releasing inflammatory cytokines^7^; and (3) reactive astrocytes form an encapsulating sheath around the electrodes in the following weeks, increasing impedance as neuronal loss ensues within the recordable radius of the injury site^7^. This last phase, in which astrocyte function plays a complex role in the immune response and device performance, remains poorly understood: how does reactive gliosis contribute to suboptimal performance? Is it merely a barrier to signal transmission and a source of increased impedance, or does it influence the function of local neurons^9^? Recent literature suggests that reactive astrocytes are highly heterogeneous, and may have either beneficial or detrimental effects on the healing of central nervous system damage^10^. These effects are injury dependent, meaning their expression is determined by the detected change in the brain microenvironment following insult. Ischemia tends to lead to the activation of signal transducer and activator of transcription 3 (STAT3), which induces a neuroprotective state, while mechanical damage tends to lead to the activation of nuclear factor kappa B (NF-kB), which releases neurotoxic factors^11^. Neurotoxic reactive astrocytes may kill healthy neurons, and these astrocytes also are associated with a loss in the ability to maintain and support synapses^12^.

We hypothesized that reactive astrocytes may participate in the neuronal loss and synaptic changes that have been observed surrounding implanted electrodes^13,14^, potentially contributing to decreased chronic signal-to-noise ratio (SNR) for implanted electrodes. Reactive astrocytes deviate from the normal gene expression and function of brain homeostasis, and specific patterns of gene expression have been associated with neurotoxic or neuroprotective astrocytic phenotypes^11^. Changes in gene expression have been noted surrounding devices in recent literature by identifying differentially expressed (DE) genes of astroglial scarring post explantation^15^; these DE genes include many of the biomarkers associated with neurotoxic astrocytes induced by inflammation^15,16^. Recent work has expanded on this body of knowledge by comparing the change in genetic expression of implanted and naïve tissue at varying distances away from the injury site, for different timepoints^16,17^. However, the nature of the transcriptional profile of reactive astrocytes encapsulating implanted devices remains unclear.

We sought to investigate the relationship between gene expression in glial fibrillary acidic protein (GFAP)-expressing tissue surrounding silicon arrays arrays to unmask biomarkers of device-reactive astrocytes. To do this, we explored the similarity between cultured reactive astrocyte models and the transcriptional profiles associated with astroglial scarring surrounding implanted electrodes in the rat brain, as revealed via combined immunohistochemistry and spatial transcriptomics. We investigated two reactivity models: rat cortical astrocytes exposed to either microglial-derived cytokines^12^, or lipopolysaccharide (LPS, a more generalized reactivity model that induces an infection-like response)^18^. We found that the astrocytes surrounding devices bore a more similar expression pattern to cytokine-induced astrocytes than LPS-exposed cells. By investigating the expression pattern between device-associated astrocytes and cultured models, we revealed novel genes associated with the tissue response surrounding devices, including genes associated with lipid metabolism and neurodegeneration. These results revealed new insight into the phenotype of device-reactive astrocytes, as well as the utility of *in vitro* models to explore reactivity *in vivo*.

## Materials and Methods

### 2.1 Cell Culture

Our cytokine-induced cell culture model was inspired by the reactive astrocyte model that was used by *Liddelow et al*.^12^, which was derived from methods developed by *Foo et al*.^19^. In the Foo and Liddelow methods, they subjected their postnatal astrocytes to several rounds of immunopanning to purify their cultures. To simplify our approach, and considering the heterogeneity expected *in vivo*, we did not immunopan our cultures. As discussed previously^20^, astrocyte maturity is an important consideration when inducing a reactive state. While we used embryonic (E-18) rat astrocyte cells, we passaged them for a total of four times over ∼4 weeks of continuous culture prior to ∼1 week of serum withdrawal and treatment; literature suggests that embryonic glial cultures have the characteristics of mature glia after they have been in culture for roughly 35 days^20^.

For our study, E-18 embryonic rat cortical astrocytes (Gibco, 4×10^6^) were seeded in a 6-well plate at a cell density of 200,000/cm^2^ in Dulbecco’s Modified Eagle Media (DMEM, Thermo Fisher) supplemented with 15% fetal bovine serum (Gibco) for a total number of four passages (1:3 split per passage). Our pilot experiments using qPCR indicated that *C3*, a biomarker for cytokine-induced reactivity^12^, was induced most reliably following four passages *in vitro* (∼4 weeks in culture, data not shown). Cells were then transferred into serum-free media for a period of six days prior to 24 hours of treatment with a cytokine cocktail or LPS. As such, the entire culture period is comparable to the 35 day time frame recommended to achieve adequate glial maturity^20^. More specifically, on the fourth passage, astrocytes were transferred to 24-well plates and treated with a serum-free media containing 50% neurobasal media (Gibco, 21103-049), 50% DMEM (DMEM, Invitrogen, 11960-044), 292 µg/mL L-glutamine (Invitrogen, 25030-081), 1mM sodium pyruvate (Invitrogen, 11360-070), 5µg/mL N-acetyl cysteine (NAC, Sigma, A8199), 100 U/mL penicillin, and 100µg/mL streptomyocin (Invitrogen, 15140-122), as described by previous protocols^12^. Every 3 days, the cells were supplemented with 5 ng/Ml HBEGF (Peprotech, 100-47), for a total incubation time of 6 days *in vitro*, and then treated for 24 h with C1q (30 ng/mL, MyBioSource, MBS143105), IL-1α (3 ng/mL, Sigma I3901) and TNF (30 ng/mL, Cell Signaling Technology, 8902SF), based on reported methods^12^. Separate serum free astrocytes were treated with lipopolysaccharide (LPS, O55:B5, Sigma Aldrich, 1µg/mL) for 24 h based on reported methods^18^. Control cells were treated identically, but without added LPS or cytokine exposure. Following the 24 hour period, media was removed and cultures were thoroughly rinsed with PBS. After the buffer was completely aspirated, plates were stored at - 80C until RNA extraction. Reported data were produced from samples pooled from two separate, repeated culture experiments (*n*=4 wells/condition for each experiment).

### 2.2 RNA Extraction and Sequencing

Total RNA was extracted from cultured reactive astrocytes using RNAzol (Molecular Research Center, Inc) or RNEasy (Qiagen) extraction kits according to the manufacturers’ instructions. The samples were then submitted to the University of Michigan Advanced Genomics core for library preparation and sequencing. Samples were subjected to 150 base paired end cycles on the NovaSeq-6000 platform (Illumina). Differential expression analysis was performed by the University of Michigan Bioinformatics Core. Data were first pre-filtered to remove genes with 0 counts in all samples. Differential gene expression analysis was performed using DESeq2^21^, using a negative binomial generalized linear model (thresholds: linear fold change >1.5 or <-1.5, Benjamini-Hochberg FDR (Padj) <0.05). Plots were generated using variations of DESeq2 plotting functions and other packages with R version 3.6.3. Genes were annotated with NCBI Entrez GeneIDs and text descriptions. GraphPad Prism was used to generate plots (DotMatics).

### 2.3 Spatial Transcriptomics

Tissue samples analyzed in this study were included in a previous report^17^. Data was gathered using methods previously described^17,22^. Briefly, a 10x Genomics (Pleasanton, CA) spatial gene expression platform (“Visium”) was used to assess brain tissue sections from Sprague-Dawley rats implanted with a silicon Michigan-style electrode (A1×16-3mm-100-703-CMLP, 15 um thickness, NeuroNexus Inc, AnnArbor, MI) for either 24 hours, 1 week, or 6 weeks (1 rat per time point). All animal procedures were approved by the Michigan State University Animal Care and Use Committee. Each spot with spatially resolved gene expression in Figures 3-6 is 55µm in diameter and 100µm apart, center-to-center. Prior to sequencing, tissue sections were labeled using immunohistochemical methods for neuronal nuclei (rabbit anti-NeuN, 1:100, Abcam, cat. #: 104225) and glial fibrillary acidic protein (mouse anti-GFAP, 1:400, Sigma-Aldrich, cat. #: G3893), and cell nuclei were counterstained using Hoechst (10 µg/mL). Images were collected as previously described^17,22^. Using the LoupeBrowser software interface (10X Genomics), differential gene expression analysis compared the gene expression of the cluster of spots within 150µm of the device interface to the cluster of spots greater than 500µm from the device interface, excluding the spots under the glia limitans. As an exception, differential gene expression in the 24-hour spatial transcriptomics sample was calculated by comparing the whole tissue section to a naïve, non-implanted tissue section. This discrepancy is due to previous observations of broad (>500µm) DE gene expression at the 24 hour time point^17^. This yielded 5811 differentially-expressed (DE) genes at 24 hours, 1056 DE genes at 1 week, and 163 DE genes at 6 weeks (p<0.05).

### 2.4 Comparison of RNA-sequencing of Cultured Reactive Astrocytes to Spatial Transcriptomics

DE genes from each timepoint *in vivo* were compared to DE genes from the *in vitro* cytokine and LPS-treated astrocyte models. Genes were selected for further investigation if they appeared in both the *in vivo* and *in vitro* experiments and their LFC had the same sign (positive or negative). Lists of genes meeting these criteria were made for each *in vivo* timepoint for both the cytokine and LPS *in vitro* conditions (24 hours/cytokine, 24 hours/LPS, 1 week/cytokine, 1 week/LPS, 6 weeks/cytokine, 6 weeks/LPS). Complete lists can be found as Supplementary Tables 1-3.

A limitation of the Visium assay is its spatial resolution: with a 55µm diameter spot size, the currently available technology does not reach single-cell resolution. For further analysis of cell type-specificity, we employed a matrix-factorization strategy using custom code in MATLAB (MathWorks, Natick, MA). In this approach, the counts of each gene detected in each spot are assumed to be derived from the mixture of the individual cell types present at each spot. In order to infer the cell types present at each spot, we used a list of 40 genes associated with 4 major cell types in the central nervous system (neurons, astrocytes, microglia, and oligodendrocyte precursor cells, OPCs)^23^. Because it is a gene with known association with the astroglial response to implanted electrodes, we included GFAP in our astrocyte gene list. For each cell type, we used low-rank Non-negative Matrix Factorization (NMF)^24–26^ to factorize the data into spatial cell counts and cell-type expression profiles.

Prior to importing the Visium 10x datasets into MATLAB, the data were first loaded into the Seurat R toolkit. In Seurat, the data was normalized using SCTransform^27^. After normalization, the data is exported into MATLAB and NMF is applied. The NMF is optimized using the Alternating Direction of Multipliers Method (ADMM)^28,29^.

From preliminary calculations (rank = 1,2,3), we find the following best model ranks:

Astrocytes: 1

Neurons: 2 (two neuron profiles distinguished mainly by expression of Sst, Stmn2, and Tmem130)

OPCs: 1

Microglia: 1

Adding additional rank to the model then gives only small changes in gene expression or spatial cell counts, rather than revealing new categories of data.

After learning the spatial cell count map for each cell type, the similarity between reactivity gene spatial maps and cell-type maps was computed. Similarity between the two maps was defined as cos(*θ*), where *θ* is the relative angle between the spatial maps when viewed as vectors where each spatial point is a dimension, based on the formula v1*v2 = |v1||v2|cos(*θ*) (Supplementary Table 4). The cell type bearing the greatest similarity in marker expression to the spatial map of the gene of interest was considered to contribute maximally to the expression pattern of that gene. Similarly, the cell type with the least similarity in marker expression to the spatial map of the gene of interest was considered to have the least contribution to the expression pattern of that gene. In this way, we could provide an initial assessment of the relationship between reactivity genes detected in cultured astrocytes and specific cell types in the spatial transcriptomics samples. For comparison, we also used Seurat to estimate Astrocyte density, making use of the Allen Institute mouse cell-type dataset^30–33^. We find in comparison to the NMF model trained on the normalized data, that the Seurat cell density underweights the evidence of Astrocyte presence provided by *Gfap* counts (Supplementary Fig. 1). A visual comparison of the spatial distribution of NMF-identified cell types and reactivity markers can be found in Supplementary Fig. 2.

**Figure. 1.**
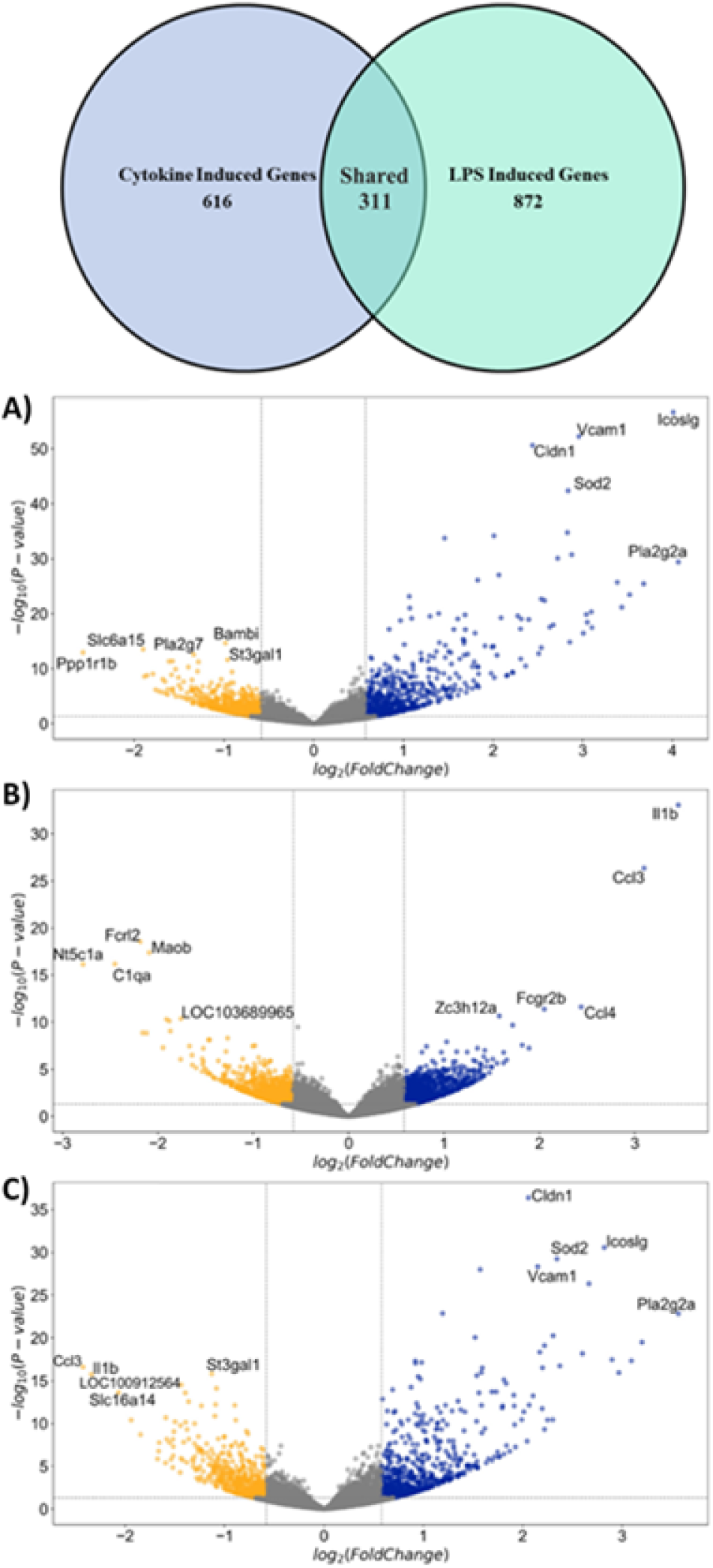
Volcano plots of LI’S and cytokine-induced reactive astrocyte models. Both cytokine treatment and LPS exposure induced hundreds of significantly DE genes in comparison to control cells, where 311 genes were shared between the two models (Venn diagram above). (A) Cytokine vs. Control, (B) LPS vs. Control, and (C) Cytokine vs. LPS. Both cytokine treatment and LPS exposure induced hundreds of significantly DE genes in reference to control cells. Significance was determined as a Log_2_FoldChange > 0.6. and p_adj_< 0.05.

## 3. Results

### 3.1 Differential Gene Expression in Cell Culture Models

We detected 927 differentially expressed genes between cytokine-treated cells and controls, 1,183 DE genes between LPS-treated cells and controls, and 831 genes between cytokine- and LPS-treated cells (Fig. 1). We noted that certain genes were significantly upregulated in both models (“pan-reactive” genes) and others were specifically upregulated only in cytokine-induced (IL-1α, TNF, and C1q) or LPS-treated cells (“unique” genes). Of the significantly DE genes, 311 were shared between the two models (Fig. 1), yielding 616 unique cytokine-induced genes and 872 unique LPS-treated genes. Complete differential expression results, as well as spatial transcriptomics data, are supplied as supplementary files.

#### 3.1.1 Unique Cytokine-Induced Genes

As expected, cytokine treatment induced upregulation of complement-associated genes. The complement cascade is involved in the innate immune response and includes an important class of genes upregulated in both neurodegenerative diseases^10,12^ and traumatic brain injury (TBI)^34^. Complement component 3 (*C3*), which was identified as a key biomarker of cytokine-induced astrocytes in the seminal report by Liddelow *et al*.^12^, was significantly and specifically upregulated in cytokine-treated cells in our data (Fig. 2). Likewise, complement factor B (*Cfb*) upregulation was unique to our cytokine-induced culture model. *Cfb* encodes an astrocyte-derived factor that supports survival of microglia^35^, which promotes synaptic pruning in the phagocytic reactive state.

We also noted the increased expression of enzymes associated with the reactive phenotype conversion in cytokine-induced cells. Phospholipase A2 (*Pla2g2a*) registered the highest fold change of any gene, and it was uniquely DE in the cytokine condition. Phospholipase A2 is an enzyme that hydrolyses phosphoglycerides to yield fatty acids and leads to the production of eicosanoids^36^. *Pla2g2a* was found to be expressed in reactive astroglia only in the areas where neuronal death occurred^36,37^. Other enzymes detected in our data are associated with the transition to a reactive phenotype, which occurs when astrocytes sense and react to exogenous agonists^38–40^ and neurotransmitters released from presynaptic clefts^38,41,42^. Astrocytic end-feet express G-protein coupled receptors (GPCRs) that become phosphorylated by Rho GTPase 1, which is the enzyme that becomes activated when activating microenvironment changes occur. We observed increased expression of Rho family GTPase 1 (*Rnd1*) specifically in the cytokine-induced astrocytes. Phosphorylation of GPCRs in astrocytes encourages calcium channel expression^38^, and literature suggests an upregulation in calcium channel expression occurs during the conversion of normal to reactive astrocytes^18,43^. Another upregulated enzyme, Receptor Interacting Serine/Threonine Kinase 2 (*RIPk2*), contains a C-terminal caspase activation and recruitment domain (CARD) and is a component of signaling complexes in both the innate and adaptive immune pathways^44^. It is an upstream regulator and potent activator of NF-kappaB and is an inducer of apoptosis in response to various stimuli^44^. NF-kappaB activation is required for neurotoxic reactive astrocyte expression^11^. A downstream biomarker, superoxide dismutase 2 (*Sod2*), is a free radical scavenging enzyme which is expressed in Alzheimer’s Disease, aging, ischemic stroke, and Parkinson’s Disease^45^. Interestingly, it also has been identified in explanted rat tissue surrounding the injury site of microelectrode array shanks^46^. Taken together, the induction of these enzymes following cytokine exposure aligns with the conversion of these cells to a reactive state.

**Figure. 2.**
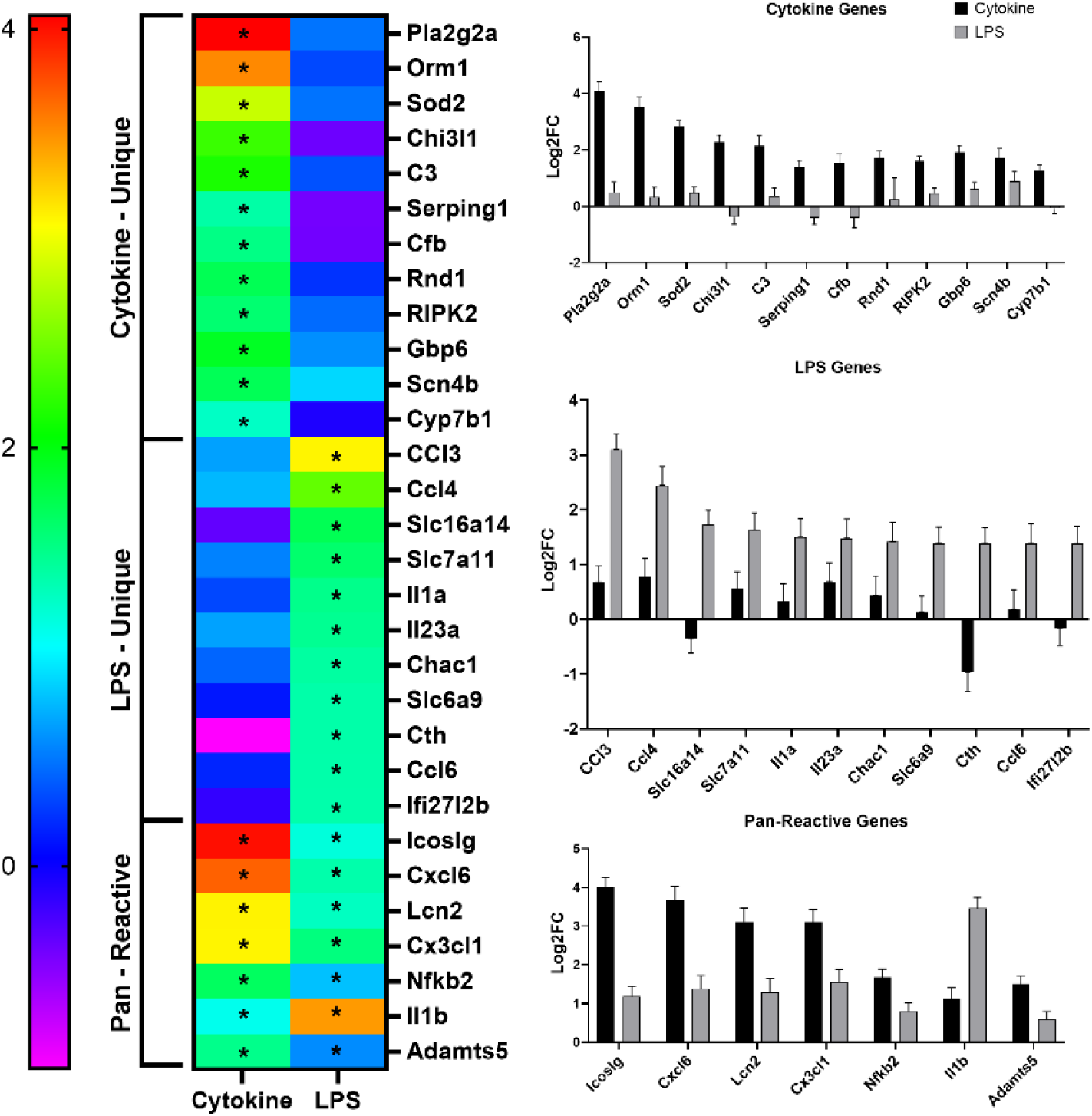
Comparison of LPS and cytokine-induced reactive astrocyte models. A selected subset of examples of individual genes which were significantly differentially expressed in each condition, as well as those which were shared (“pan-reactive”), are illustrated in the heatmap. Color bar shows Log_2_FoldChange the versus control cells, and “*” denotes statistical significance (p_adj_<0.05). On the right, individual bar graphs illustrate specific genes and their associated Log_2_FoldChange for each condition. Error bars represent the standard error of the Log_2_FoldChange.

We also observed an upregulation of the expression of several genes associated with ion channel expression and activation. Guanylate binding protein family member 6 (*Gbp6*) is a protein, induced by interferon, which hydrolyzes guanine triphosphate (GTP) to guanine diphosphate (GDP) and guanine monophosphate (GMP). It is catalyzed by Rho GTPase and plays a role in calcium channel activation in astrocytes. Another detected gene, sodium voltage-gated channel beta subunit 4 (*Scn4b*), activates Na_v_1.5 in the wake of mechanical damage. *Scn4b* is specifically linked to [Ca^2+^]_i_ gradient shifts and promotes glial scarring^47^. Changes in ion channel expression have been observed surrounding devices in previously reported data, where Na_v_ expression developed an inverse relationship with K_v_ expression during the progression from acute to chronic injury^48^. More recently, we observed a cluster of ion channel-associated genes upregulated in rat brain tissue as a part of the acute-phase injury to electrode insertion in motor cortex^17^.

#### 3.1.2 Unique LPS-Induced Genes

LPS activates the Toll-like receptor 4 (TLR4)-dependent pathway that induces microglia to release IL-1β, IL-6 and TNFα via the mitogen-activated protein kinase (MAPK) and NFkB pathways^49^. As expected, IL-1α and IL-23α were observed as uniquely upregulated in our LPS-treated cells. There was a lesser association of this model with complement-associated genes, which is not unexpected because the complement component system is activated by pathogens, whereas LPS is an endotoxin. True to what is observed in reactive astrocyte transcriptomes, we also observed the upregulation of chemokine genes in LPS-treated cells. This included the upregulation of inflammation-associated factors such as C-C motif chemokine ligand 3 (*Ccl3*), which is induced typically through the STAT3 pathway, and is reduced through nicotinamide adenine dinucleotide phosphate oxidase (NOX2) inhibition^50^. Chen *et al*.’s studies suggest that NOX2 inhibition may provide neuroprotection against inflammatory damage^50^. These observations align with the view that a reactive astrocyte population exists on a spectrum of reactivity; some may be neurotoxic and some may be neuroprotective^10^. In addition, C-C motif chemokine ligand 4 (*Ccl4*) is upregulated in the LPS condition and has been associated with neurodegenerative diseases including Alzheimer’s disease and HIV-associated dementia^51^. Finally, C-C motif chemokine ligand 6 (*Ccl6*) was upregulated in LPS-treated cells. *Ccl6* has been associated with mediation of astroglial migration^52^. Its expression in chronic reactive astrocytic populations surrounding explants provides evidence of reactive astrocytic migration^52,53^. All three of these chemokines (*Ccl3, Ccl4, Ccl6*), which were selectively upregulated in our LPS-treated astrocyte cultures, are associated with oxidative stress-induced brain injury^46,50–52^.

Endotoxins can cause apoptosis, and we observed enzymatic biomarkers that are key regulators of apoptotic pathways.Glutathione Specific Gamma-Glutamylcyclotransferase 1 (*Chac1*) has been shown to promote neuronal differentiation by deglycination of the Notch receptor, ultimately inhibiting neurogenesis^54^. This enzyme depletes glutathione, which is a necessary factor in the apoptotic cascade^54^. Solute carrier family 6 member 9 (*Slc6a9*) plays a role in shuttling glycine during deglycination and has been found to be upregulated in patients with other neurodegenerative diseases associated with erythrocyte dysregulation, such as cerebral malaria^55^. Additionally, there are genes in our data which are linked to alternative pathways to arrive to apoptosis. For example, a common way to tag proteins for cell death is through methylation. Cystathionine gamma-lyase, (*Cth*), which is upregulated in our data, does this by converting cystathionine to cysteine. This increases the elevation of total homocyesteine (tHcy), a risk factor expressed in Alzheimer’s disease and cognitive loss^56^.

In a broader sense, we observed differentially expressed genes in our data set that are associated with neurodegenerative disease states, and those biomarkers can provide context for the nature of astrogliosis^57^. Interferon, alpha-inducible protein 27 like 2B (*Ifi27l2b*), that participates in apoptotic signaling, has been observed in our LPS dataset and also in *Zamanian et al*.*’s* reactive astrocyte transcriptome data^53^. Another biomarker observed in our LPS data set which is associated with disease states, such as Alzheimer’s disease and cognitive loss, is solute carrier family 16, member 14 (*Slc16a14*). *Slc16a14* codes for monocarboxylic acid transporters, which are highly expressed in the hippocampus and hypothalamus^58^.Likewise, solute carrier family 7 member 11 (*Slc7a11*), is highly expressed in glioma patients and may be responsible for seizures^59^; it is upregulated in microenvironments experiencing neurotoxic levels of glutamate^59^.

### 3.2 Comparisons of Gene Induction in Cell Culture Models to the In Vivo Tissue Response

To contextualize the cellular origin of recently observed changes in gene expression surrounding electrodes, we sought to compare gene expression in cultured astrocyte models to gene expression surrounding devices implanted in the brain. To do this, DE genes from each timepoint *in vivo* were compared to DE genes from the *in vitro* cytokine and LPS-treated astrocyte models.

Genes were selected for further investigation if they appeared in both the *in vivo* and *in vitro* experiments and their LFC had the same sign (positive or negative). Lists of genes meeting these criteria were made for each *in vivo* timepoint for both the cytokine and LPS *in vitro* conditions (24 hours/cytokine, 24 hours/LPS, 1 week/cytokine, 1 week/LPS, 6 weeks/cytokine, 6 weeks/LPS). Complete lists can be found in Supplementary Table 5. Cytokine and LPS intersections with the 24 hour *in vivo* DE genes each yielded 256 and 260 genes, respectively. The Cytokine and LPS intersections with 1 week *in vivo* DE genes yielded 104 and 66 genes, respectively. The Cytokine and LPS intersections with 6 week *in vivo* DE genes yielded 38 and 21 genes. Further analysis separated pan-reactive genes (those common to all models) from genes unique to each reactivity model to determine which reactive astrocyte model, Cytokine or LPS, is most aligned with *in vivo* observations. After pan-reactive genes were removed from the *in vivo* vs. cytokine intersections, there were 165 unique genes found at 24 hours, 71 genes found at 1 week, and 23 genes found at 6 weeks. After pan-reactive genes were removed from the *in vivo* vs. LPS intersections, there were 169 unique genes found at 24 hours, 33 genes found at 1 week, and 6 genes found at 6 weeks. The composite expression patterns of these genes, at each time point, are shown in Figures 3 and 4 (these figures illustrate genes with positive LFCs; negative LFCs can be found in Supplementary Fig. 3). While gene expression changes were broadly spatially distributed at the 24 time point, the cytokine and LPS associated genes were more clearly colocalized with GFAP-expressing scar tissue at later time points (insets, Fig. 4).

**Figure 3.**
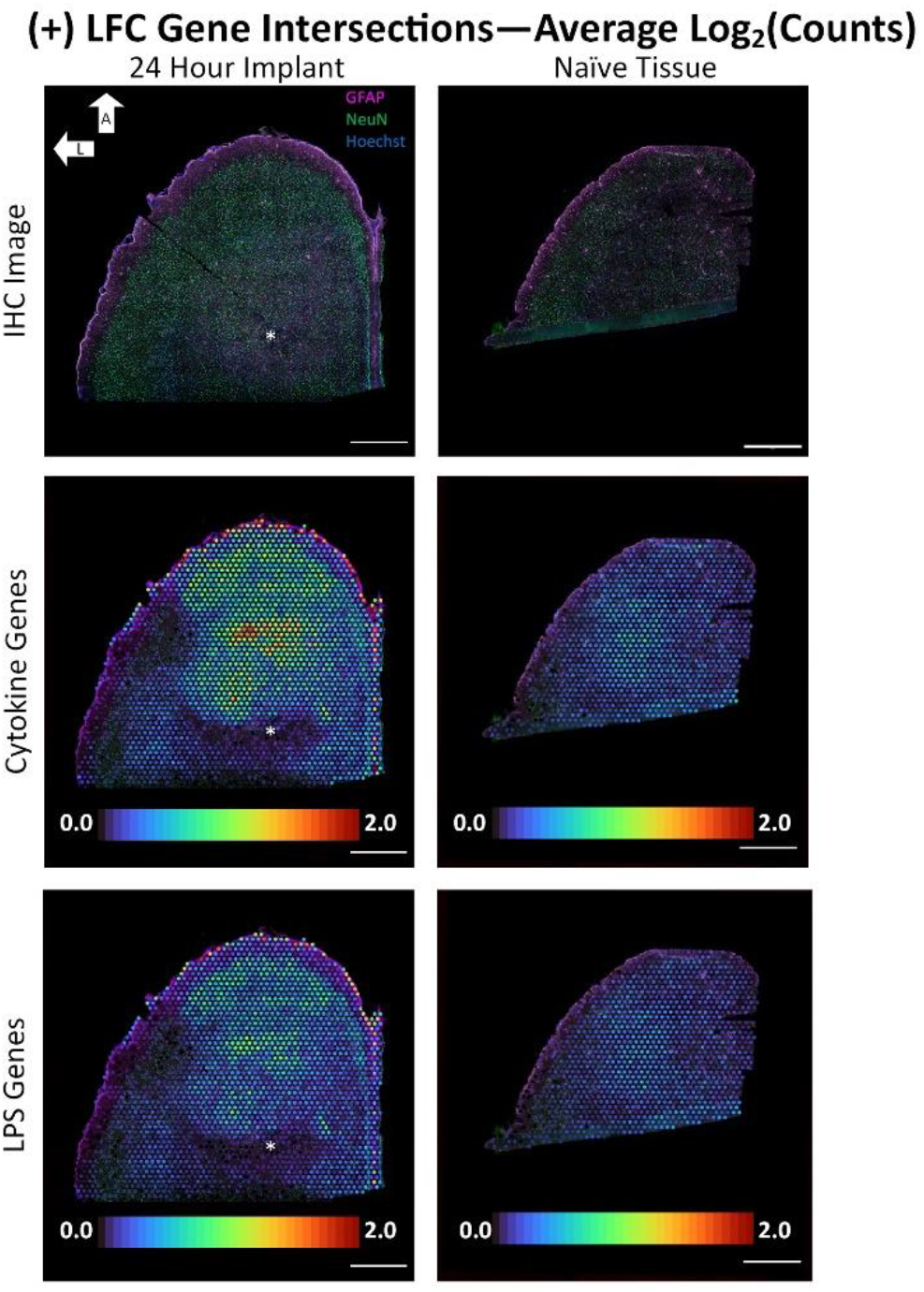
Average expression of genes differentially expressed both in astrocytes *in vitro* and in a 24-hour electrode implant *in vivo*. IHC images show a transverse tissue section of brain tissue implanted for 24 hours (left) and an unplanted, naïve brain tissue section (right). Purple: GFAP, Green: NeuN, and Blue: Hoechst; L: Lateral and A: Anterior. Cytokine panels show the average number of counts for the genes that were found to be positively differentially expressed in both the cytokine treated astrocyte cultures and in a 24-hour electrode implant *in vivo* compared to the naïve tissue section. LPS panels show this same data except for genes differentially expressed in the LPS treated astrocytes and the 24-hour implant. Scale bars: 1000µm.

**Figure 4.**
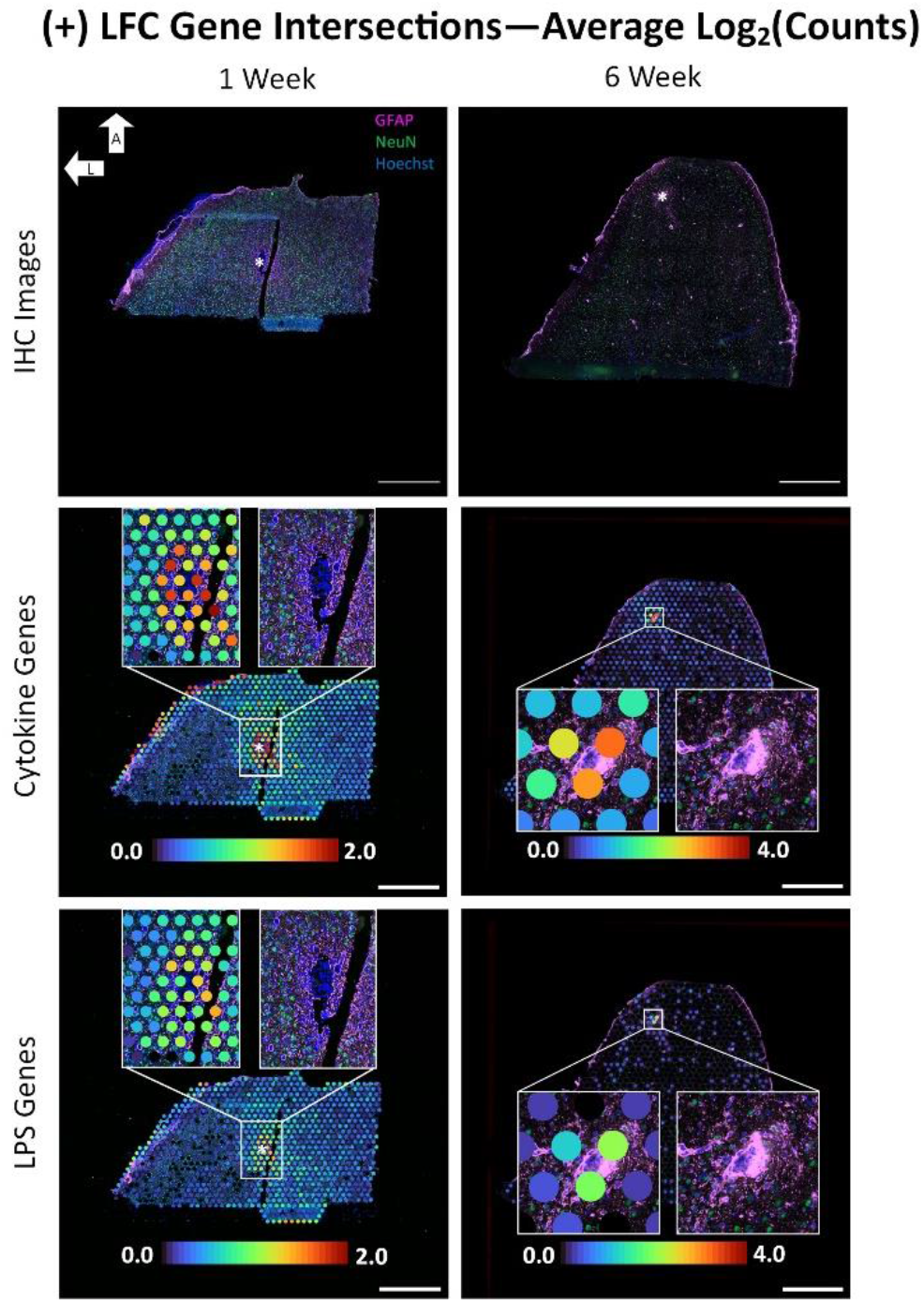
Average expression of genes differentially expressed both in astrocytes *in vitro* and in 1-& 6-week electrode implants *in vivo*. IHC images show a transverse tissue section of brain tissue implanted for 1 week (left) and 6 weeks (right). Purple: GFAP, Green: NeuN, and Blue: Hoechst; L: Lateral and A: Anterior. Cytokine panels show the average number of counts for the genes that were found to be positively differentially expressed in both the cytokine treated astrocyte cultures and in *in vivo* tissue sections. LPS panels show this same data except for genes differentially expressed in the LPS treated astrocytes and the 1-and 6-week implants. Scale bars: 1000µm.

We further explored the spatial patterns of the expression of individual cytokine-induced genes surrounding the device tracts at each time point. Examples of individual genes at each time point are displayed in Figures 5 and 6. These genes were selected for display based on a combination of high differential expression in the cytokine-induced culture model, localized expression associated with the electrode implant, and relevancy to reactive astrocytes based on reported literature. The cytokine-induced genes included significant differential expression of both complement component 3 (*C3*) and *Serping1*, as reported previously^12^. Upon inspection of the electrode interface, we observed tight clustering of the expression of these genes in the compact GFAP-expressing scar surrounding the device tract (Fig. 5). As a complement-associated protein, astrocytic C3 expression is associated with the neuroinflammatory response in neurological injuries and diseases, as well as with the loss of synaptic connectivity between neurons^12^. Recent work reported elevated *C3* expression within 100 microns of the electrode interface^16^, and the present study revealed overlap of *C3* expression with the presence of GFAP-positive astrocytes surrounding the electrode.

**Figure 5.**
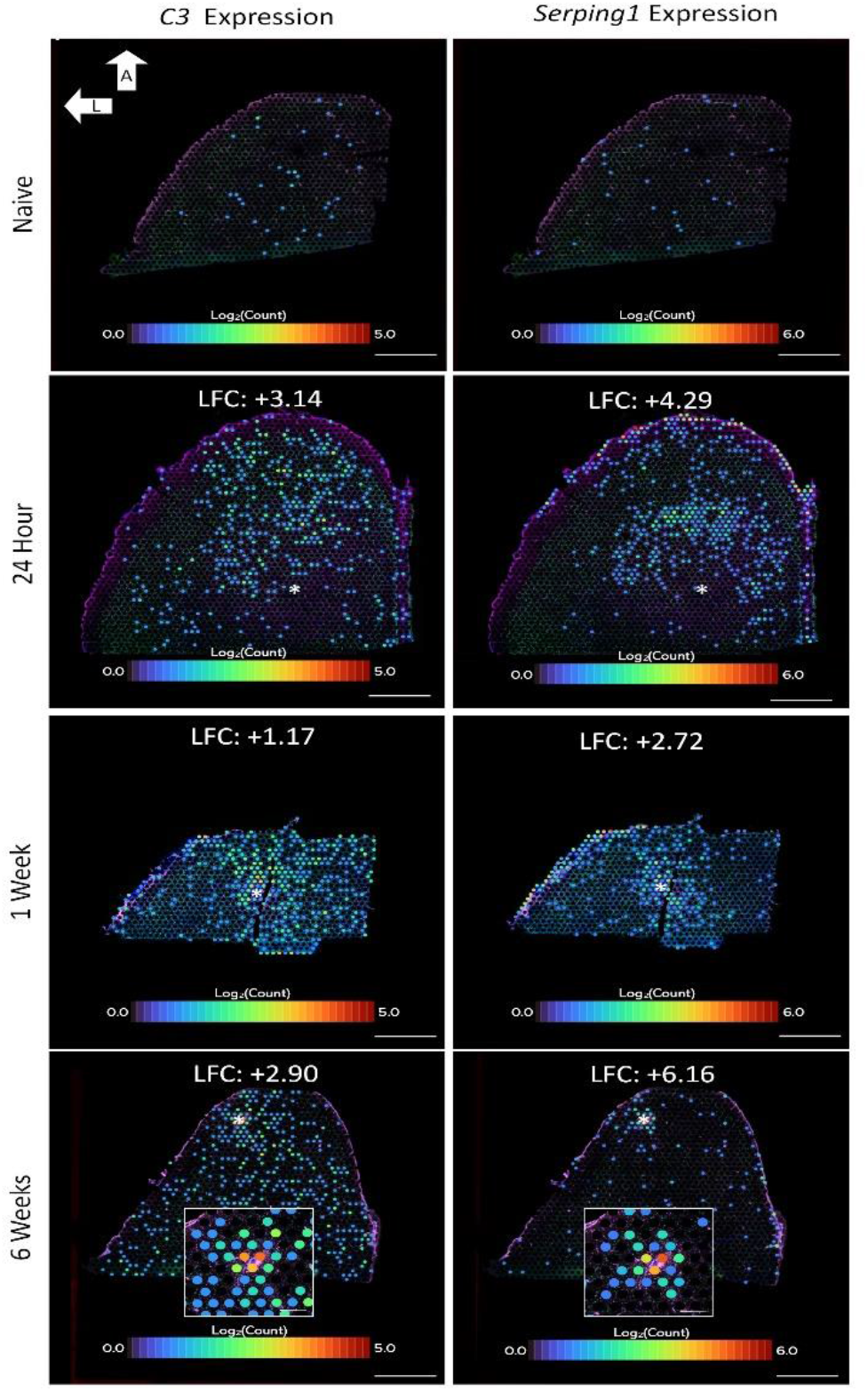
Individual expression of reactive astrocyte genes *C3* and *Serping1*. **(Left Column)** Log_2_(counts) of *C3* at each implant timepoint as well as in an unimplanted, naïve control. Expression of *C3* is increased in each of the implant samples compared to the naïve sample, with the exact LFCs shown above the image of each sample. **(Right Column)** Log_2_(counts) of *Serping1* at each implant timepoint as well as in an unimplanted, naïve control. Expression of *Serping1* is also increased in each of the implant samples compared to the naïve sample, with the exact LFCs shown above each image. L: Lateral, A: Anterior. Scale bars: 1000µm.

**Figure 6.**
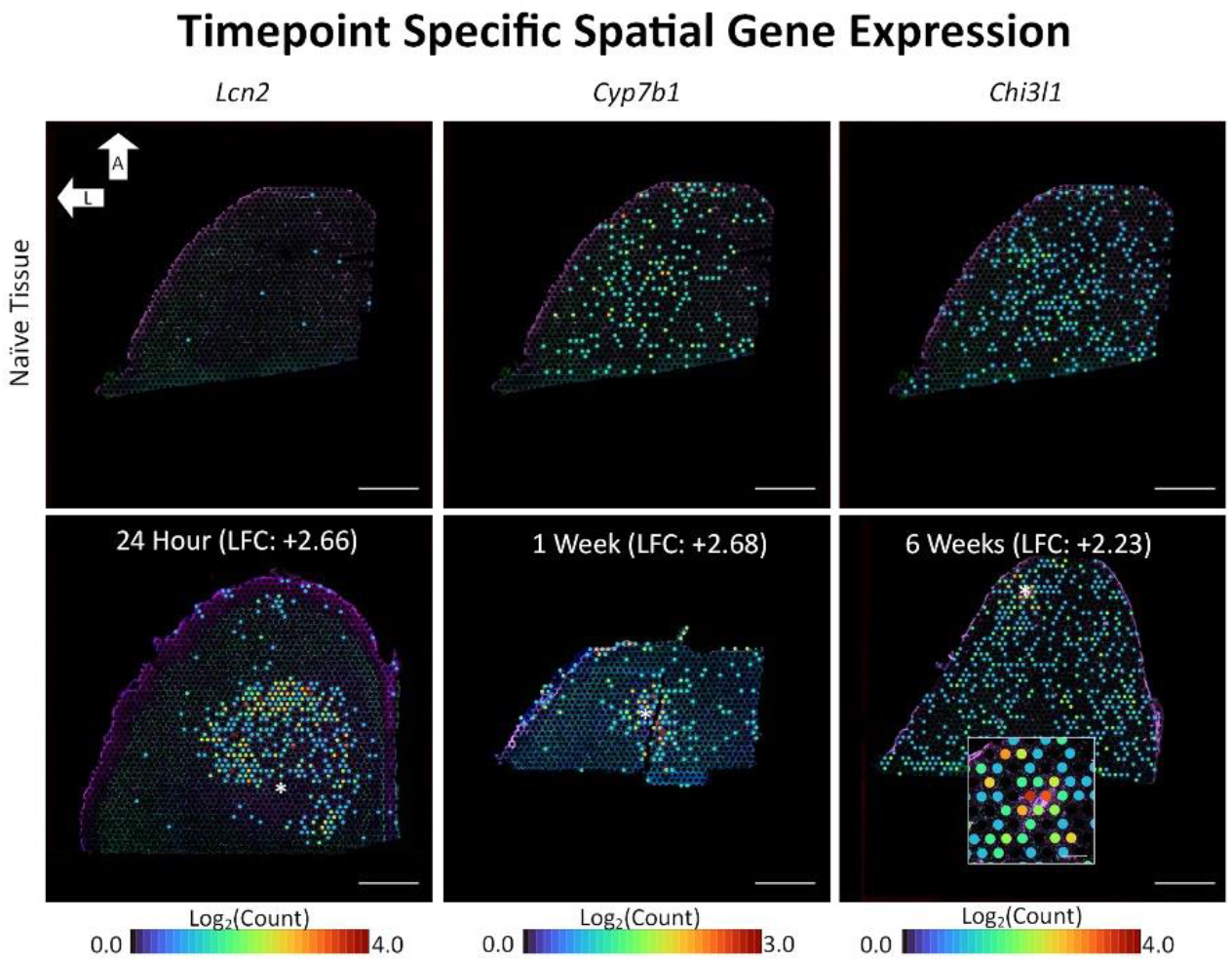
Individual expression of timepoint-dependent genes of interest. *Left Column*. Log_2_(counts) of *Lcn2* in the 24-Hour implant sample (bottom panel) compared to its expression in naïve tissue (top panel). *Middle Column*. Log_2_(counts) of *Cyp7b1* in the 1-week implant sample (bottom panel) compared to its expression in naïve tissue (top panel). *Right Column*. Log_2_(counts) of *Chi3l1* in the 6-week implant (bottom panel) compared to its expression in naïve tissue (top panel). LFCs for each gene are shown above the implant image for each timepoint and are compared to that gene’s expression in the naïve sample. L: Lateral, A: Anterior. Scale bars: 1000µm.

In addition to these effects, which were forecasted by previous transcriptomic profiling in cytokine-induced astrocytes, we observed additional markers of device-reactive astrocytes through our comparative investigation (Fig. 6). For example, at the 24 hour time point *in vivo, Lcn2* was found upregulated and was also upregulated in the cytokine *in vitro* dataset. *Lcn2* is a potent neurotoxic protein that can be secreted by astrocytes^60^. At the 1 week time point, cytochrome P450 family 7 subfamily B member 1 (*Cyp7b1*) was noted. Cyp7b1 is an enzyme needed for the synthesis of an oxysterol implicated in astrocyte migration^61^. Chitinase 3-like protein 1 (*Chi3l1*) was revealed in astrocytes at the device interface at the 6 week time point^62^. *Chi3l1* encodes a secreted glycoprotein (YKL-40) which has been associated with reactive astrocytes and neurodegeneration in several reports^62,63^. Its expression is known to increase in parallel with tau in Alzheimer’s disease (AD), and a recent investigation in a mouse model of AD suggested that, “*Chi3l1* may suppress glial phagocytic activation and promote amyloid accumulation^62^.” It is possible that its expression at the 6 week time point surrounding electrodes could serve to dampen and constrain glial reactivity as the device interface stabilizes. Since tau pathology has been observed at chronic time points (16 weeks) surrounding implanted electrodes, it is likewise possible that *Chi3l1* contributes to emergence of AD-associated markers surrounding implants^64^. Future work will need to be conducted to explore the relationship between *Cyp7b1* expression and the formation of an astroglial sheath, as well as a subsequent, potentially multi-faceted role for *Chi3l1* in constraining glial responses at the expense of neuronal health. These are two examples of genes revealed through our analysis; raw data files and analysis results are provided for further investigation (see Supplementary data).

The spatial resolution of the transcriptomics assay used in this study is improved in comparison to previous methods used at the implanted electrode interface^16^, yet it does not reach single-cell resolution. We used a factorization strategy to add insight into the relationship between reactivity genes highlighted in our results (*C3, Serping1, Cyp7b1*, and *Chi3l1*) and the distribution of astrocytes in the associated 1-week and 6-week spatial transcriptomics samples. We found that *Serping1, Cyp7b1*, and *Chi3l1* were most strongly correlated with the presence of astrocytes while *C3* was most strongly correlated with the presence of microglia (Supplementary Table 4). These associations were consistent across all three time points.

## 4. Discussion

In this study, we compared reactive astrocyte models to the foreign body response to implantable recording neurotechnology. The novel potential biomarkers identified could uncover knowledge about the brain’s foreign body response and reveal new targets for treatment. Likewise, we hope to develop a practically useful *in vitro* model to study the foreign body response, and eventually, enable high throughput screening. An *in vitro* testbed would be valuable to test hypotheses and develop approaches to screen new electrode designs and intervention strategies more efficiently, and gene expression is a potentially useful readout to enable the development of such an assay. Previous studies have sought to identify culture conditions which model the foreign body response to electrodes implanted in the brain. An early iteration of this approach employed a ‘scrape’ injury, or the placement of a segment of wire, within a co-culture of microglia, astrocytes, and neurons^65^. Observations of glial migration and device encapsulation recreated familiar elements of device-tissue interaction. Several subsequent reports iterated on this approach, with improvements including the addition of oligodendrocytes^20^, the inclusion of inflammatory mediators such as lipopolysaccharide^65^, and the development new culture protocols to reduce the experimental burden associated with long, continuous cultures^20^.

Our approach focused specifically on astrocyte reactivity as a potential pathway of interest, with the goal of informing recent observations of device-induced changes in gene expression using spatial transcriptomics^16^. Our methods were derived from studies using LPS to induce inflammation in astrocytes^18^, as well as a recently reported, cytokine-induced astrocyte model^12^. The latter approach is of particular interest, as the microglial source of cytokine release, as well as reported effects on synaptic connectivity, are reminiscent of the brain tissue response to electrodes. While each culture model expressed genes relevant to inflammatory responses at the device interface, our inspection indicated a closer correspondence of the cytokine-induced model with the *in vivo* tissue response (Fig. 4). Markers associated with “neurotoxic” astrocytes that lose their ability to support the maintenance of neuronal synapses were upregulated surrounding implant sites (*C3, Serping1*; Fig. 5)^12^. Both evidence of a reduction in excitatory synapses surrounding implants, as well as the observation of increased *C3*, have been observed previously at the device interface^14,16,66^. Comparing *in vivo* gene expression data with the transcriptional profiles of cultured reactive astrocyte models is a potential path toward revealing the mechanisms initiating reactivity in device-responding cells. Additionally, we propose that spatial transcriptomics data may be a viable path toward optimizing culture-based models of device-tissue interactions: by iteratively benchmarking *in vitro* results relative to *in vivo* data, it may be possible to identify the most accurate culture models by comparing transcriptional profiles between the two experimental paradigms.

The upregulation of complement genes is an aspect of the cytokine-induced astrocyte model which resembles *in vivo* effects. However, these device-associated effects also may be related to microglial activation, which could induce astroglial activation and synaptic degradation. In order to assess the relationship between reactivity markers and specific cell types, we implemented a factorization strategy to determine the similarity between spatial maps of reactivity genes and cell type specific markers. Our results implicated astrocytes as the strongest contributor to the expression of our selected reactivity genes of interest (*Serping1, Chi3l1, and Cyp7b1*). Alternatively, *C3* expression was most strongly associated with the presence of microglia. The latter effect is likely intertwined with the astrocytic response, since increased expression of C3 in microglia may be related to the induction of a reactive state in neighboring astrocytes via cytokine release^12^. While our analysis represents an initial attempt to associate *in vitro* transcriptional profiles with specific cell types *in vivo*, the results suggest that reactivity genes detected in the astrocyte culture models were strongly associated with astrocytic presence at the device interface.

It is important to acknowledge that limitations remain in our current study: our simplified culture method may have increased astrocytic heterogeneity, and questions remain regarding the similarity of these models to responses in the intact brain. While our current assessments relied on gene expression, future analysis may include detailed morphometric analysis, which could compare changes in astrocytic structure associated with reactive states in the *in vitro* and *in vivo* settings. Additional opportunities for future work include using more advanced computational techniques, which may elucidate biomarkers more effectively. Armed with an increasing knowledge base of the relationship between devices and gene expression^15–17^, it may be possible to further refine culture models to more faithfully recreate these responses in future work. It may also be possible to directly test the impact of identified biomarkers on electrode function by implementing strategies to manipulate gene expression at the device interface^67^.

It also remains to be determined how translatable the current results will be to the clinical setting. Rodents currently are the gold standard *in vivo* model used to study the tissue response to implants. However, compared to the intact human brain, rodent models have less diverse astroglial populations, limiting the ability to accurately predict human patient outcomes^68^. Also, because of a short life cycle, murine models are not reliable indicators of potential long-term performance in human patients. Additionally, 2-dimensional culture models suppress important genetic expression^69^ because they do not actively mimic the biochemical and mechanical properties of the 3-dimensional brain microenvironment. The dynamic organelle that houses the brain microenvironment, the extracellular matrix (ECM), consists of proteoglycans and glycosamioglycans such as hyaluronic acid, meaning the mechanical properties of the functional tissue are governed by these macromolecules^68^. Astroglial cells have mechanosensing abilities, and placing these cells in a dish results in the diminishment of important homeostatic cell signalling.

Emerging techniques using human induced pluripotent stem cell (iPSC)-derived, 3-dimensional brain organoids^70,71^ may address many of the limitations associated with murine models and 2D cultures. Spatial and organizational support is needed for self-assembly to support constant rearrangement and separation of cells, as observed in development during the creation of a structurally robust ECM. Patterned protocols mimic specific areas of the brain while integrated protocols make whole brain organoids^71^ to produce vasculature and immune type cells through co-culturing methods^72–75^. In the future, human iPSC brain organoids may be an interesting model to study DE genes surrounding electrodes implanted in the brain. In addition to recreating the more complex environment of intact brain tissue, it may also be possible to test the tissue response to electrodes in the context of models of neurodegenerative disease. In this study, we have reported the differential expression of genes that have been associated with multiple sclerosis and Alzheimer’s disease. Differential expression of disease-associated genes around implanted devices raises questions regarding whether implanted electrodes have the potential to exacerbate the expression of these genes in subjects which may be susceptible or predisposed to developing a neurodegenerative disease. Upregulation of genes associated with accumulation of amyloid protein such as *Chi3l1* and other potential AD genes (*Cth, Slc16a14, Sod2*) could suggest a possible interaction between implanted technologies and disease markers. The growing use of transcriptomics in the study of device-tissue interaction already has opened up new understanding of the biological mechanisms of the tissue response to implanted electrodes [15-16,22,73]^15–17,22,76^, paving the way for the future extension of these methods to investigate effects in clinically relevant models of human disease.

## Acknowledgements

The authors thank: Rebecca Tagett and the Bioinformatics Core of the University of Michigan Medical School’s Biomedical Research Core Facilities for RNAseq analysis support; Dr. Steven Suhr and Dr. Marie-Claude Senut of Biomilab, LLC, for assistance with early experiments and RNA extraction; Samuel Daniels for contributions to initial pilot experiments; and Dr. Mark Reimers for consultation on methods for inferring the cell type specificity of results. This funding was supported by the National Institutes of Health (R01NS107451 and F99NS118738).

## Data Availability Statement

Raw data is available upon request to the corresponding author.

## Notes

### Competing Interest Statement

The authors have declared no competing interest.

