## Supplementary Table 4 for "Gene Expression Changes in Cultured Reactive Rat Astrocyte Models and Comparison to Device-Associated Effects in the Brain"

|  | **C3** | **Serping1** | **Chi3l1** | **Cyp7b1** | **GFAP** |
| --- | --- | --- | --- | --- | --- |
| **Astrocyte** | 0.34679 | 0.49499 | 0.78872 | 0.36487 | 0.98738 |
| **Neuron 1** | 0.33502 | 0.27227 | 0.67713 | 0.25264 | 0.7799 |
| **Neuron 2** | 0.19092 | 0.16924 | 0.47467 | 0.18247 | 0.53537 |
| **OPC** | 0.35499 | 0.30767 | 0.68115 | 0.26981 | 0.78863 |
| **Microglia** | 0.3912 | 0.42477 | 0.46476 | 0.29382 | 0.60699 |

**24 Hour**

**1 Week**

**6 Weeks**

|  | **C3** | **Serping1** | **Chi3l1** | **Cyp7b1** | **GFAP** |
| --- | --- | --- | --- | --- | --- |
| **Astrocyte** | 0.50332 | 0.36858 | 0.58872 | 0.57007 | 0.77919 |
| **Neuron 1** | 0.49233 | 0.23127 | 0.55735 | 0.50935 | 0.55214 |
| **Neuron 2** | 0.40507 | 0.18819 | 0.4578 | 0.42066 | 0.46901 |
| **OPC** | 0.49891 | 0.2342 | 0.55521 | 0.50908 | 0.55217 |
| **Microglia** | 0.50385 | 0.36003 | 0.52782 | 0.46614 | 0.612 |

|  | **C3** | **Serping1** | **Chi3l1** | **Cyp7b1** | **GFAP** |
| --- | --- | --- | --- | --- | --- |
| **Astrocyte** | 0.57659 | 0.62048 | 0.66404 | 0.49183 | 0.9772 |
| **Neuron 1** | 0.38018 | 0.2153 | 0.3797 | 0.20968 | 0.52508 |
| **Neuron 2** | 0.53047 | 0.32016 | 0.50246 | 0.32039 | 0.71981 |
| **OPC** | 0.53149 | 0.31557 | 0.49185 | 0.29871 | 0.70292 |
| **Microglia** | 0.59127 | 0.61202 | 0.56111 | 0.46066 | 0.83697 |

**Supplementary Table 4.** Similarity between the reactivity and cell type spatial maps was defined as cos(*θ*), where *θ* is the relative angle between the spatial maps when viewed as vectors where each spatial point is a dimension, based on the formula v1*v2 = |v1||v2|cos(*θ*). Larger values of cos(*θ*) indicate stronger similarity between the two maps, and thus, a stronger association between the cell type and reactivity marker. Maximum and minimum values are highlighted in red and blue, respectively.
