## Supplementary Figures for "Gene Expression Changes in Cultured Reactive Rat Astrocyte Models and Comparison to Device-Associated Effects in the Brain"

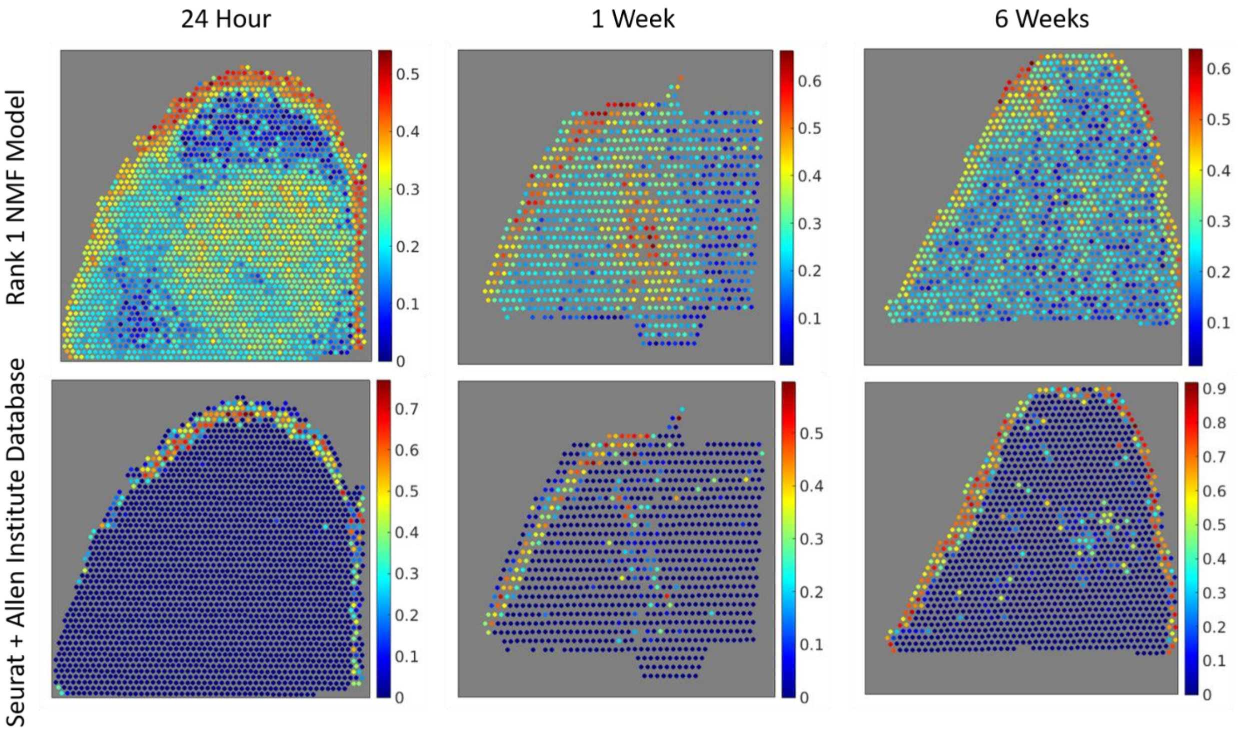


**Supplementary Figure 1.** Visual comparison between the Rank 1 NMF model (including GFAP as an astrocytic marker) and Seurat model indicate that the NMF model more faithfully detects astrocytic expression in accordance with GFAP immunohistochemistry in Visium samples (see main text figures).

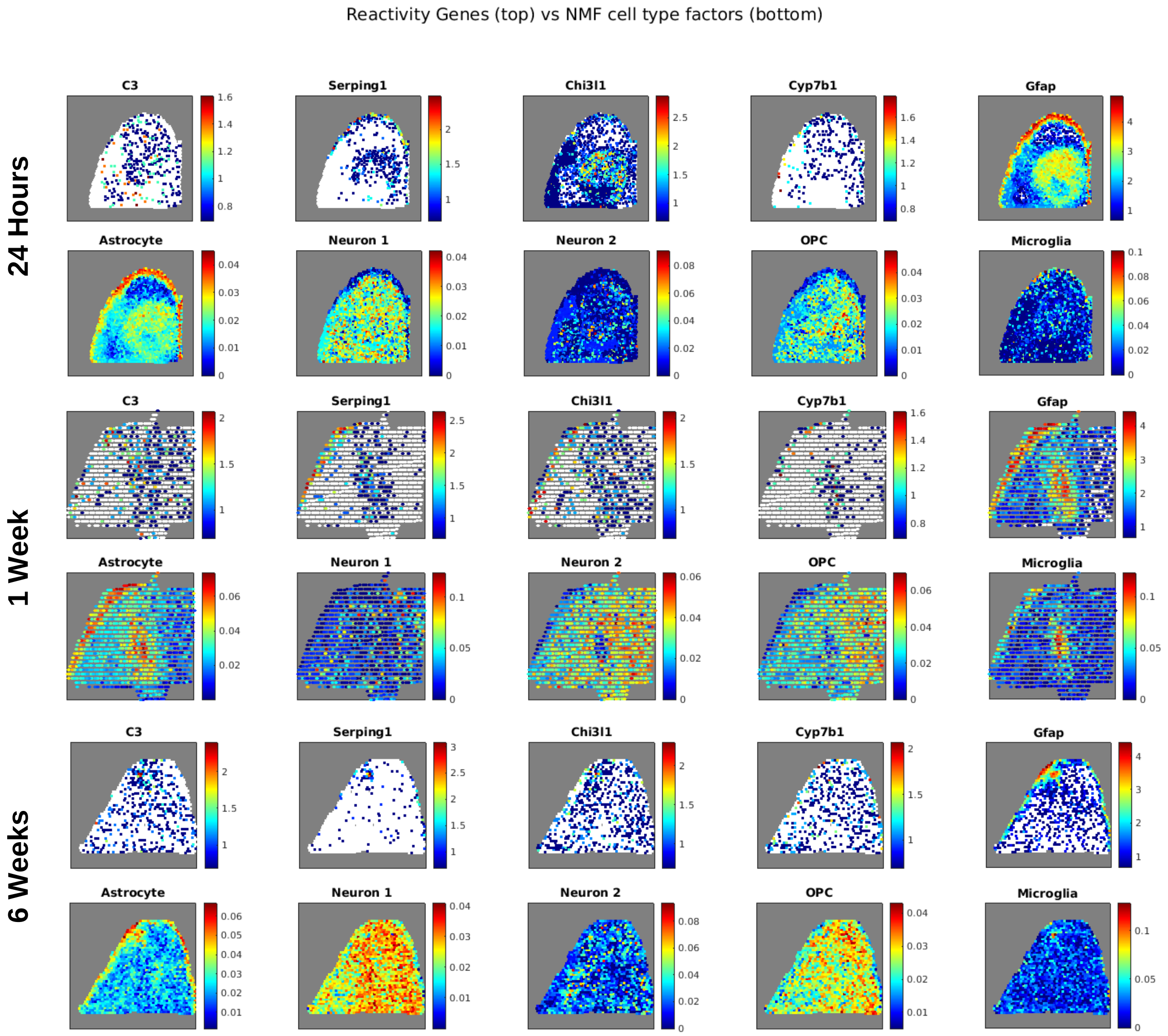


**Supplementary Figure 2.** Visual comparison between the expression pattern of reactivity genes of interest with the Rank 1 NMF model (including GFAP as an astrocytic marker) for the spatial distribution of each cell type.

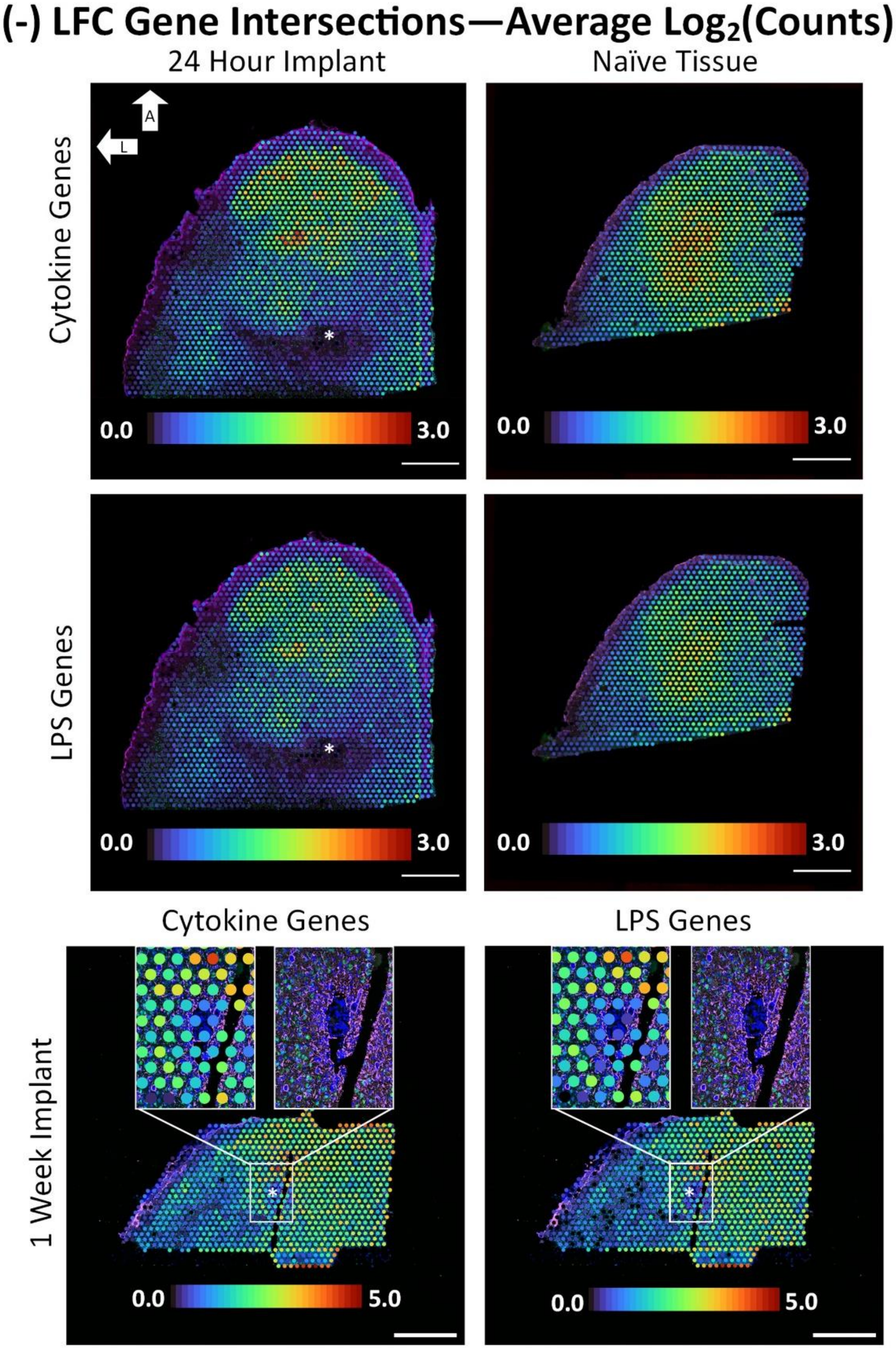


**Supplementary Figure 3.** Average expression of negative LFC genes differentially expressed both in astrocytes *in vitro* and surrounding electrode implants *in vivo*.
